# Using Metacognitive Strategies to Improve Academic Performance in Biochemistry

**DOI:** 10.1101/2020.07.08.193649

**Authors:** Heather B. Miller, Melissa C. Srougi

## Abstract

Growing evidence suggests that students’ self-beliefs about the ability to alter their academic abilities can directly influence long-term achievement. These self-beliefs or mindsets can either be fixed (unchangeable) or growth oriented. Students with growth mindsets believe their academic abilities can change, which leads to higher grades and academic persistence in contrast to students with fixed mindsets. However, less is known about how these attributes affect student learning, particularly in college level biochemistry courses. In this study, we utilized social-psychological interventions to promote growth mindset among third and fourth year undergraduate students enrolled in a one semester Biochemistry survey course. Using a mixed-methods study design we evaluated student mindset, attitudes towards learning, and academic performance over four semesters. Our results suggest that although students’ mindsets did not change as a result of metacognitive interventions, their positive perceptions about learning vs performance did increase. Furthermore, students receiving growth mindset interventions significantly outperformed students who did not receive interventions on the final cumulative exam that assessed critical thinking skills. These results suggest that metacognitive interventions can be an effective tool to improve student academic performance in a biochemistry course.

## Introduction

It is increasingly evident that students’ psychology or academic mindsets have a critical role in educational achievement. Social-psychological interventions alter ways in which students perceive themselves and help them pursue activities/opportunities that will promote academic growth. These interventions have been used widely to promote academic success across K-12 classrooms [1, 2]. One social-psychological intervention in particular focuses on the perception of intelligence. Students’ mindsets (a.k.a. implicit theories) are their own self-beliefs about personal characteristics, such as intelligence. Some individuals view intelligence as unchangeable (i.e. fixed mindset) whereas others view intelligence as malleable and that it can be changed over time (i.e. growth mindset) [3, 4]. Since students with growth mindsets tend to pursue challenges, learn, and develop their abilities despite challenges (as opposed to students with fixed mindsets), it is not surprising that growth minded students exhibit greater learning and achievement from the time they are in elementary school through college. Fostering growth mindsets can improve students’ motivation, academic achievement, and reduce racial, gender, and social class gaps. Growth mindset interventions can be disparate, but all emphasize student self-reflection and are mindful of students’ subjective experiences within an educational context [2]. Previous research has demonstrated that fostering a growth mindset can benefit underperforming students, underrepresented minorities and women in math and science [5-7], thus narrowing achievement gaps. In the transition to secondary schools, 20% of United States students will not finish high school on time [8]. However, growth mindset interventions used with these students showed improved grades, especially for those with lower achievement [9]. Growth mindset interventions have been implemented in courses to varying degrees of effectiveness, depending upon students’ implicit theories of intelligence and baseline discipline specific-skills [10]. The most powerful positive effects have been demonstrated during the transitions into junior high and high school [6, 7, 11-13].

While research on metacognition has largely been focused on K-12 students [14], more recent studies have examined its effects in post-secondary students in the humanities and social sciences [14-17]. However, the use of metacognitive interventions (although assumed to be effective across disciplines) has not been fully investigated in biochemistry classrooms. Two independent groups reported using metacognitive interventions with students enrolled in a general chemistry I and introductory biology course, respectively, demonstrated a selective effect on underrepresented minority students by reducing the achievement gap [18, 19].

The present study investigates undergraduate Biochemistry students’ self-reported mindset beliefs and their implications for students’ academic performance. Our research questions were 1) would growth mindset interventions promote a growth mindset in biochemistry students and 2) would they increase student learning outcomes. Indeed, studies in other STEM disciplines have suggested such strategies are useful in improving student academic performance [18, 20]. To test this, we designed and implemented a variety of metacognitive interventions in a one semester Biochemistry course. The series of metacognitive interventions aimed to alter students’ perceptions of academic challenges commonly encountered in upper-level interdisciplinary courses. Biochemistry courses in particular challenge students to use higher order thinking to integrate knowledge from a variety of previous course work (e.g. biology, general, and organic chemistry). This study evaluates the mindset and academic performance of undergraduate students enrolled in a Biochemistry course who were exposed to metacognitive interventions compared to student groups who were not exposed. Our results suggest that metacognitive interventions can improve student academic performance in Biochemistry. Furthermore, these interventions can be readily implemented into a variety of courses at both the small and large scale and do not require specialized technology or monetary resources.

## Methodology

### Student Demographics

Students enrolled in this study were 18 years or older and enrolled in Biochemistry I during 4 consecutive fall semesters at High Point University. This lecture course was designed for non-biochemistry majors and there was no corequisite lab required. The prerequisites for the course were a freshman level biology course, two semesters of general chemistry, and two semesters of organic chemistry. Students participating in this course were from numerous majors with the largest enrollment from chemistry, exercise science and biological science majors. Students in the non-intervention group were composed of a total of 74 undergraduate students (juniors and seniors) with a 34% male to 65% female ratio. Students in the intervention group were 87 total undergraduate students with a 35% male to 64% female ratio. Participation was completely optional, and students completed consent forms during the first week of class. Incentive to participate consisted of one bonus point on the final exam. Those who did not consent to the study were given an alternative bonus question equal in weight. The High Point University Institutional Review Board (IRB) approved this protocol (#201607-499) under an exempt review.

### Instructor Information

Two instructors were involved in this study, and each taught the course for two fall semesters (Fall 2015 and Fall 2016 or Fall 2017 and Fall 2018). These instructors were both tenure-track faculty and members of the Chemistry Department with similar educational/training backgrounds including doctoral degrees and post-doctoral training. Additionally, they had similar years of undergraduate teaching experience.

### Course Synopsis

“Biochemistry I” is a semester-long course covering the chemical and physical properties of proteins, nucleic acids, carbohydrates, and lipids. It also introduces students to the study of bioenergetics and carbohydrate metabolism. The course met twice a week for a total of three hours.

### Student Learning Outcomes

Course expectations and assessment methods were detailed in the syllabus, as were the learning outcomes:

Upon completion of this course, students should be able to:

1. Explain the chemical and physical properties of each type of macromolecule, enzymes, carbohydrates and nucleic acids.
2. Describe protein-based information technologies, signaling, energetics and biochemical reactions in a biological context.
3. Explain the steps of glycolysis, gluconeogenesis, TCA cycle, and oxidative phosphorylation.
4. Interpret data from primary scientific results and peer reviewed journal articles.

### Assessment Methods

Student learning outcomes were measured by a variety of assessment methods throughout the semester. At the beginning and end of the course, students’ foundational biochemistry knowledge was assessed by a validated Biochemistry Diagnostic Assessment Instrument [21]. Students were given the same assessment instrument at the conclusion of the course to assess learning gains throughout the semester. This diagnostic assessment was not used in course grade calculations, but the outcomes were shared with the students. Additionally, students were given four written exams constituting 58% of the overall grade in the non-growth mindset (non-GM) group and 55.5% of the overall grade in the growth mindset (GM) group. Other assessment methods included online quizzes (non-GM 12.5%; GM 10%) and homework (10%) as well as written reflections for the GM group (10%). Written reflections were assessed based on thoughtfulness of reflection, the ability to follow directions and proper spelling/grammar. In lieu of written reflection assignments, non-GM students were assessed on their class participation and citizenship (5%). A final comprehensive and cumulative American Chemical Society Core Biochemistry Exam was given at the conclusion of the course and counted towards 14.5% of students’ overall grades in both the non-GM and GM sections. Statistical significance of examination scores between semesters was calculated using a two-tailed Student’s *t*-test.

### Curricular Interventions

Two groups of students who enrolled during Fall 2016 and Fall 2017 were subject and participated in GM interventions (see below) that employed numerous social-psychological interventions to help them internalize and make cognitively available the belief that intelligence is malleable. The attitudes and student achievements of these students were compared to those of two control groups, Fall 2015 and Fall 2018, that were not subject and did not participate in the growth mindset (non-GM) interventions. The course content was identical in either the GM or non-GM semesters. The only difference consisted of implementation/participation of GM interventions, including the addition of reflective assignments in the GM cohort. At the start and end of the semester, participants completed an anonymous online survey to measure their beliefs about intelligence to determine their growth mindset index and their attitudes towards learning [4].

To foster growth in academic mindsets, participants enrolled in the GM sections were exposed to several interventions interwoven throughout the course structure which were: self-reflection exercises [22] concept mapping [23, 24], growth minded messaging [6, 11] exam wrappers [25, 26] and instructor talk [27]. Detailed descriptions on metacognitive interventions can be found in the Supporting Information (Appendix S1).

#### Survey

Consenting participants in the study were given time during class to complete an anonymous online survey at both the start and end of the semester. The survey used was an adapted version of the implicit theories of intelligence scale for adults to evaluate students on their mindset [4]. Mindset scores were used to calculate growth-mindset index (GMI) as described elsewhere [4, 5, 28]. Questions pertaining to attitudes towards student learning and demographic information were also included; surveys were administered and analyzed using Qualtrics software (www.Qualtrics.com, Provo, UT) and Microsoft Excel. At the completion of each survey, students were given a link to a second survey where names were entered to confirm completion. The names were not tied to any individual survey responses, and enabled instructors to award completion credit (i.e. bonus point on an in-class exam) for completion of both surveys. In the control sections, 39 and 38 students completed the pre-survey and post-survey, respectively. In the growth-mindset experimental sections, 88 and 89 completed the pre and post-survey, respectively. Chi-square analyses were performed in Microsoft Excel to calculate p-values for survey responses between groups.

## Results

### Course context

Students participating in this course were from numerous majors including biology, exercise science, neuroscience, psychology, and chemistry (**Figure 1A**). The course enrolls 30-45 students each fall and includes 75 minutes of lecture twice a week. There is not a required laboratory for this course. Instructors of the course utilized active learning techniques such as clicker questions, problem-based conceptual and mathematical questions, case-studies, and group activities. Information about students’ sex, race, major and GPA were self-reported via online survey for those providing informed consent to participate in the study. A total of 169 students enrolled in the study. Students enrolled in the course in Fall 2015 and Fall 2016 semesters were the control group and received no growth minded interventions (non-GM). In this cohort, 65.8% of respondents were female and 34.2% male; an intersex option was not available on the survey. Of the total, 92.4% (n=36) self-identified as white, whereas the underrepresented minority group made up 7.6% (n=3 and included students who were Black or African American, Hispanic or Latin@, American Indian or Alaska native, or Native Hawaiian or Pacific Islander) (**Table 1**). Students enrolled in Fall 2017 and Fall 2018 received growth minded (GM) metacognitive interventions. The GM cohort consisted of 64% female respondents and 35.9% male. Of students in the GM sections of the course, 83.3% (n=55) self-identified as white whereas 16.6% (n=6) included underrepresented minority groups (**Table 1**).

**Table 1.** Student demographic information.

| STUDENT DEMOGRAPHIC | NON-GM COHORT<br>NUMBER OF STUDENTS<br>(% TOTAL) | GM COHORT<br>NUMBER OF STUDENTS<br>(% TOTAL) |
| --- | --- | --- |
| **GENDER** |  |  |
| MALE | 13 (34.2) | 32 (35.9) |
| FEMALE | 25 (65.8) | 57 (64.0) |
| **ETHNICITY** |  |  |
| HISPANIC OR LATIN@ | 4 (10.5) | 6 (9.1) |
| NOT HISPANIC OR LATIN@ | 34 (89.5) | 60 (91.0) |
| **RACE** |  |  |
| WHITE | 36 (92.3) | 55 (83.3) |
| BLACK OR AFRICAN<br>AMERICAN | 0 (0.0) | 5 (7.6) |
| AMERICAN INDIAN OR<br>ALASKA NATIVE | 2 (5.1) | 1 (1.5) |
| ASIAN | 0 (0.0) | 2 (3.0) |
| NATIVE HAWAIIAN OR<br>PACIFIC ISLANDER | 1 (2.5) | 1 (1.5) |
| OTHER | 0 (0.0) | 2 (3.0) |

**Figure 1.**
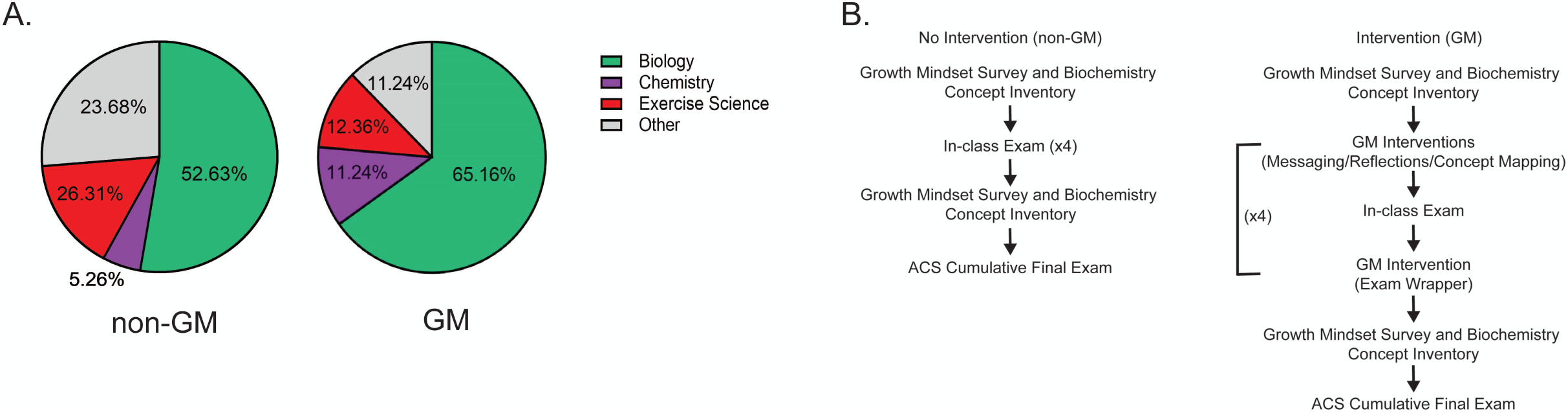
Class major and course structure. A) Student participants enrolled in non-GM and GM semesters of the study that completed the online anonymous survey were categorized based on their major. B) Diagram of course structure in which non-GM (*left panel*) or GM interventions (*right panel*) were implemented.

Students enrolled in Biochemistry I in 2016 and 2017 were exposed to practices that addressed the affective domain of learning, whereas all other course structures remained the same from the non-GM sections (**Figure 1B**). Interventions were centered around fostering an academic growth mindset. This was addressed by the consistent implementation of course activities with growth mindset messages throughout the course of the semester (**Figures S1-4**). Students were assigned six self-reflection assignments that had students reflect on challenging past experiences and how they overcame them (**Figure S1**). Instructors shared growth-minded messages by incorporating PowerPoint slides into the lecture period that highlighted the malleability of intelligence, the benefits of hard work and persistence as well as the biochemistry of adaptability and evolution (**Figure S2**). Additionally, students were required to create concept maps of key biochemistry topics to provide them with a framework to document their learning process and progress, thus emphasizing their learning gains rather than course grades per se (Example of student work, **Figure S3**). Finally, students were required to fill out exam wrappers after each in-class examination. These exam wrappers were modified to incorporate growth-minded prompts for reflection, which emphasized the importance of the learning process (**Figure S4**).

### Impact of Metacognitive Interventions on Students’ Growth Mindset and Student Attitudes of Learning

To address our first research question (determine whether students’ mindsets changed in response to growth mindset interventions), we administered an 8-item growth mindset survey at the start and end of the semester [4]. Survey results were used to determine students’ growth mindset index (GMI). GMI scores ≥ 3.5 are considered “growth minded” while scores < 3.5 are considered more “fixed minded”. The average GMI score at the start and end of the course for non-GM students was 3.29 and 2.97, respectively. The range was similar for students in the GM sections where student scores were 3.37 at the start and 3.36 at the end of the course indicating tendencies towards fixed mindedness (Figure 2A) [4]. However, there was not a statistical difference in GMI scores in student cohorts who were subject to metacognitive interventions versus students that were not subject to interventions either before (pre non-GM *vs* GM; p = 0.78) or after the course (post non-GM *vs* GM; p = 0.47) (**Figure 2A**). Interestingly, however, there was more variability in response to mindset questions in the non-GM group as well as a sharp decay in GMI at the conclusion of the course. The magnitude of loss of growth mindset thinking was attenuated in the GM group. These data suggest that metacognitive interventions may have delayed the loss of GM as seen in previous studies [29].

**Figure 2.**
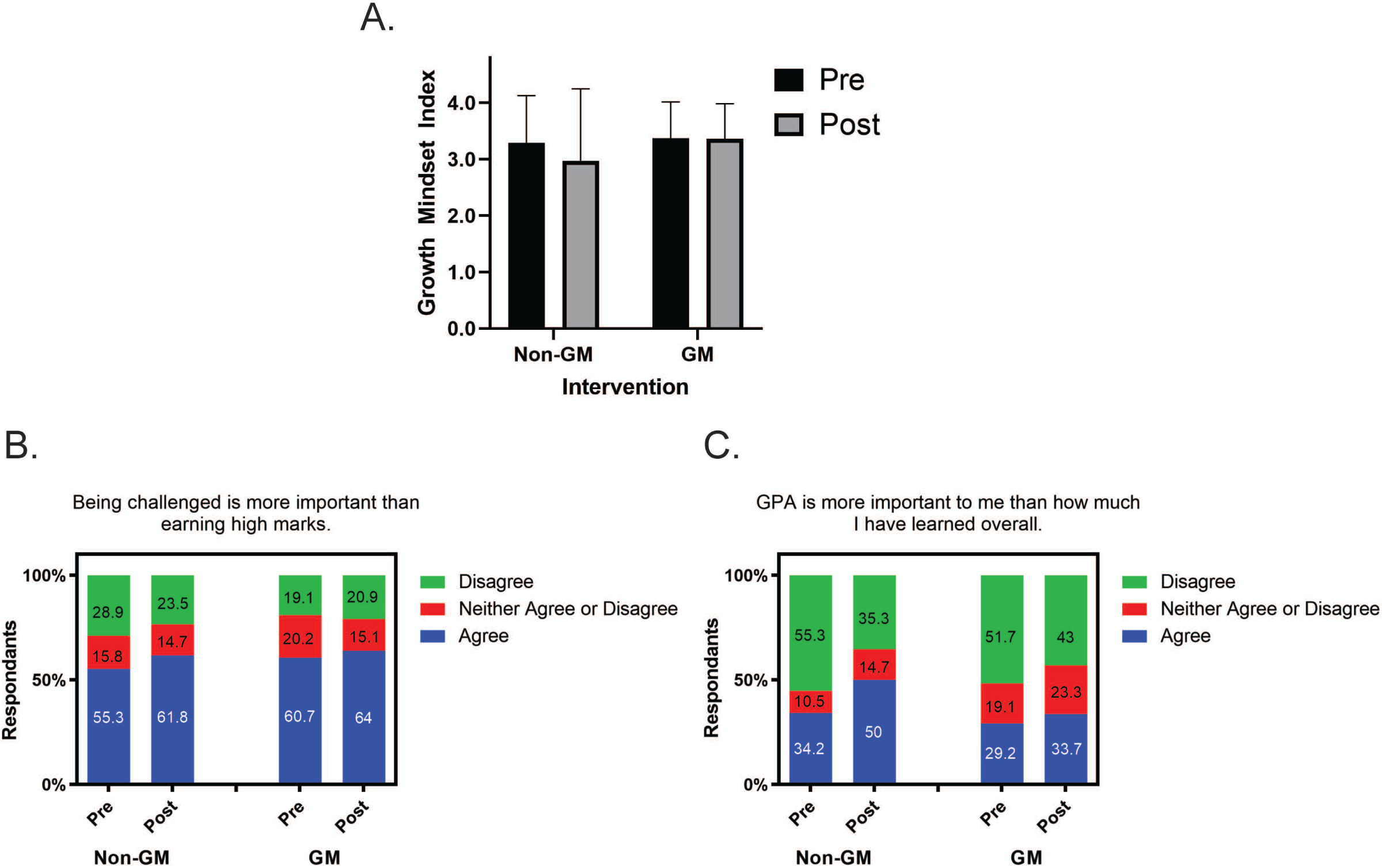
Growth mindset interventions did not alter student mindset but increased positive perceptions about learning. A) Growth mindset index was determined before (pre) and after (post) the course using the Dweck intelligence scale questionnaire. Non-GM pre *n*=39, GM pre *n*=88, (*p* = 0.78 pre non-GM *vs* pre GM) non-GM post *n*=38, GM post *n*=89 (*p* = 0.46 post non-GM *vs* post GM). Error bars represent the S.D. Statistical significance was determined using two-tailed *t*-tests. B-C) Summary of overall percentage of student responses in non-GM and GM cohorts to an anonymous 5-point Likert scale questionnaire given at the beginning (pre) and end (post) of the semester. Values indicate overall percentage responses for each specified question. B) Data showing student responses to the importance of being challenged vs earning high marks. Non-GM pre *n*=39, GM pre *n*=88, non-GM post *n*=38, GM post *n*=89. (*p* = 0.79 post non-GM *vs* post GM) C) Data showing students responses to the importance of GPA vs learning. Non-GM pre *n*=39, GM pre *n*=88, non-GM post *n*=38, GM post *n*=89. (*p* = 0.08 post non-GM *vs* post GM). Statistical significance was determined using a Chi-square analysis.

We next wanted to determine if students’ perceptions of learning were altered in response to metacognitive interventions. Student attitudes on a number of items were analyzed through an anonymous online survey administered at the beginning and end of each semester. Students with growth-minded views of intelligence value learning as a process and seek challenges [4]. These survey questions asked students their views on the importance of grades versus the learning process. Students were asked to respond to the following statement, “Being challenged is more important than earning high marks.” At the beginning of the semester, 60.7% of students in the GM group and 55.3% of students in the non-GM group agreed with this statement (pre non-GM *vs* GM; p = 0.37) (**Figure 2B**). By the end of the course, these percentages increased in each group. Sixty-four percent of GM students agreed that being challenged was more important than earning high marks, whereas 61.8% of non-GM students agreed (**Figure 2B**). The change in response to this statement was not significant (post non-GM *vs* GM; p = 0.78) but demonstrated a positive trend in student perceptions of the learning process.

We anticipated that since the majority of students enrolling in this course were in a pre-professional track (i.e. pre-medical, pre-dental, pre-physician assistant etc.), there would be a greater student perception of the importance of grade point average since pre-professional programs currently rely on GPA as one of the metrics for acceptance in these programs. We therefore asked students to respond to that statement, “GPA is more important to me than how much I have learned overall.” Twenty-nine percent of students in the GM group agreed with this statement at the beginning of the course, while 34.2% of students in the non-GM group agreed (pre non-GM *vs* GM; p = 0.22) (**Figure 2C**). However, these numbers both increased by the end of the semester, where 33.7% of GM and 50% of non-GM students now agreed GPA was more important than learning. Although both groups had a shift towards believing GPA was more important than learning, the change was not as profound in the GM group (post non-GM *vs* GM; p = 0.08) (**Figure 2C**). Taken together, these data are suggestive that GM interventions may influence student perceptions of learning.

### Impact of Metacognitive Strategies on Student Academic Performance

We also wanted to determine if metacognitive interventions improved student academic performance. To address this second research question, students were given the Biochemistry foundational concepts instrument both before and after the course to measure learning gains [21]. This instrument was administered in class without the students’ prior knowledge, and thus students did not prepare outside of their regular class preparations. Students in both groups demonstrated significant learning gains regardless of intervention (pre *vs* post non-GM; *p < 0.001 and pre *vs* post GM; **p < 0.01; **Figure 3A**). By the conclusion of the course, both non-GM and GM cohorts achieved a mean score of 10 correct biochemistry questions out of the 21 tested (**Figure 3A**) compared to 7 correct questions at the start of the semester. It is important to note that although this instrument was scored, it was not used towards course grades. Therefore, best efforts may not be represented accurately in this instrument.

**Figure 3.**
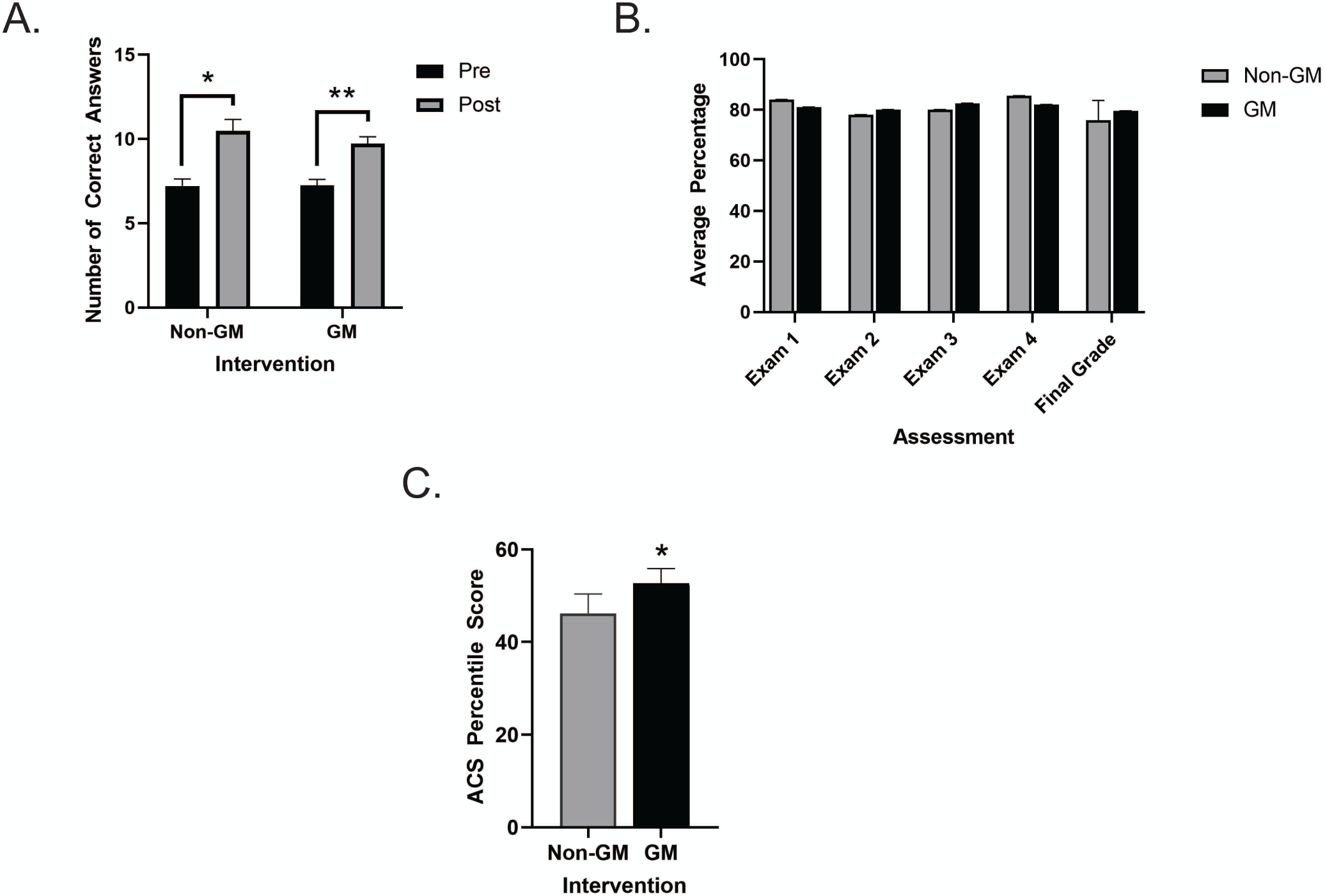
Students receiving GM interventions perform better on the final cumulative ACS Core Biochemistry exam. A) Biochemistry concept inventory assessment scores earned at the beginning (pre) and end (post) of the semester, non-GM *n* = 43, GM *n =* 88 (*t*-test \**p* < 0.001, \*\**p* < 0.01). B) Data shown are the mean in-class exam and final grades in the course for both non-GM (*n* = 71) and GM (*n* = 90) groups. C) Average percentile score on the ACS Biochemistry cumulative exam for students not receiving (non-GM *n* = 71) or receiving (GM *n* = 90) GM interventions. Error bars represent the S.E.M. Statistical significance was determined using two-tailed *t*-tests (\**p* < 0.01).

We also used additional assessment methods in the course to determine biochemistry competency, including four in-class exams given throughout the semester. There was no significant change in in-class exam scores between non-GM and GM cohorts (**Figure 3B**). The average in-class exam grade for non-GM students was an 81.8% ± 0.11 S.E.M. compared to 81.4% ± .11 S.E.M. for GM students (**Figure 3B**). At the end of the semester, students were given a standardized, cumulative American Chemical Society (ACS) Core Biochemistry final exam. This exam comprised 14.5% of students’ overall course grades. Notably, there was a statistically significant increase in academic achievement on the cumulative final exam. Students receiving metacognitive interventions significantly outperformed students that did not receive interventions, with the mean ACS percentile score of 52.7 ± 3.2 S.E.M. for GM and 46.1 ± 4.2 S.E.M. for non-GM students (p < 0.01; **Figure 3C**).

Together, these data suggest that student learning outcomes were met regardless of whether metacognitive interventions were implemented. The mean scores for the Biochemistry concept inventory and in-class exams were essentially the same for both groups. However, there was a statistically significant increase in student performance on the final cumulative national ACS exam for students who were exposed to GM interventions, suggesting that these classroom changes can have a positive impact on student academic performance.

## Discussion

The goal of this study was to experimentally test whether social-psychological interventions, such as growth-mindset interventions, can improve undergraduate biochemistry student mindset and academic performance. To do so, we first administered a variety of metacognitive interventions such as concept mapping, messaging, and self-reflection to undergraduate students enrolled in a one-semester Biochemistry survey course. We predicted that mindset interventions would benefit students particularly by shifting their mindsets from fixed- to growth-oriented [11, 30]. However, our data showed no statistically significant effect on students’ mindsets (**Figure 2A**). Meta-analysis of mindset interventions demonstrate that interventions do not necessarily boost all students’ outcomes and are most impactful for underrepresented or marginalized students [31]. Since the vast majority of students enrolled in the GM cohort of this study were white (83.3%) and female (64%) (**Table 1**) it is not uncommon for mindset-interventions to yield weak results [31, 32]. Our study did not have a large enough population of under-served or at-risk students to evaluate these effects on target groups where mindset interventions have been shown to have the most effect [6, 30, 32]. Moreover, undergraduate students’ definitions of “intelligence” are highly context specific and may vary. Some students may view “intelligence” as knowledge whereas others may view it as “abilities’’ thus not reliably measuring mindset in the appropriate context [33]. This may influence how students respond to mindset questions. Therefore, future surveys using the mindset scale will be more specific and clear about the context of the questions about intelligence.

In addition to students’ mindsets, we also asked students questions about their attitudes towards challenge and the learning process. Students who were exposed to growth-minded messaging and activities had a higher percentage of students acknowledging the benefits of challenge and the learning process. While not statistically significant, the results trended such that growth mindset interventions may help students’ approach to effort, making them more willing to participate in activities that challenge them intellectually [34].

Finally, we answered the research question if growth mindset interventions would improve academic performance. Even though students did not exhibit statistically significant changes in growth mindset in response to our interventions, there was a statistically significant increase in academic performance. Gains in learning were demonstrated by both student groups regardless of intervention (**Figure 3A-C**), suggesting that the social-psychological interventions implemented do not hinder student performance. Most notably, students’ receiving GM interventions significantly outperformed non-GM students on the final cumulative ACS biochemistry examination (**Figure 3C**). This exam particularly assesses students’ ability to retain and apply their knowledge, as well as think critically. Of all the assessments administered, this is the one that students anecdotally find to be the most challenging. Our qualitative results may suggest that students receiving growth mindset interventions are willing to approach this challenge and thus put forth more effort [34]. Moreover, although we saw no shift in students’ mindsets with GM interventions, a number of recent studies have shown the effect of instructors’ mindsets on student academic performance. Notably, students have increased academic performance in courses taught by growth-minded faculty [35, 36]. Therefore, in this study, students may perceive their instructors’ confidence in their academic abilities to improve, which can minimize stereotype threat and motivate them to actively engage in the learning process independently of shifts in their own mindsets [36].

Collectively, our qualitative and quantitative data suggest that social-psychological interventions promoting growth mindset have benefit in improving biochemistry undergraduate academic performance and attitudes towards learning. Mindset interventions are easy to implement and do not require monetary resources. Therefore, these interventions can be a useful pedagogical approach implemented in biochemistry and other courses to promote academic achievement.

## Supporting information

Supporting Information Miller and Srougi

## Acknowledgements

The authors would like to thank High Point University (HPU) students enrolled in the course for their feedback and participation. The HPU Quality Enhancement Plan SoTL Grant provided funding for this work to M.C.S. M.C.S. would also like to thank the North Carolina State University Biotechnology Program and College of Veterinary Medicine Academy of Educators for support.

## Notes

### Competing Interest Statement

The authors have declared no competing interest.

