## Supporting Information Miller and Srougi for "Using Metacognitive Strategies to Improve Academic Performance in Biochemistry"

**Supporting Information Appendix S1**

**Curricular Growth Mindset Interventions**

Self-reflections and concept mapping

As part of their course assessment, students were asked to submit six written reflections based on instructor assigned prompts. These questions sought to have students reflect on their own personal and academic journey, identify challenges that they encountered and how they overcome those challenges. An example reflective prompt, "We have discussed a number of situations whereby an organism can alter or adapt their physiology to suit new environments or purposes. For example, the Sherpa people in the Himalayas can adapt their behavior to function in low oxygen environments, or adjusting our own diet can improve cognitive function through alterations in brain biochemistry etc. For this reflective assignment, tell me **A)** how have you adapted yourself (whether it be behavior, way of thinking, etc.) to overcome an obstacle to reach a goal in your own life? **B)** How can you use (apply) this experience to improve yourself in other areas? Give a concrete example. **C)** Finally summarize how you overcame that obstacle in Part A in one BOLDED sentence." Other reflections required students to visually outline a pathway towards growth/understanding of conceptual biochemistry core concepts using concept mapping. Furthermore, other questions were designed to have students acknowledge their own adaptability and how they could apply those skills to improve themselves in other areas.

Growth mindset messaging

Specifically, this intervention consisted of 1-2 PowerPoint slides to link the day's conceptual biochemistry topic to the message that the body and brain can physically grow and adapt to different environmental cues. An example would include a situation where environmental cues could be a low oxygen environment triggering an increased blood supply to trying new study strategies that can help students achieve learning goals. This intervention was used to help students internalize the message that their physiology is capable of changing and growing to achieve their own personal goals.

Exam wrappers

Following each of the four in-class exams, students were required to complete a reflective exam wrapper. This assignment is similar to other cognitive wrappers that have been utilized in other disciplines to help improve student learning with modification [1, 2]. Prior to receiving their graded exam, students were allocated ~10 minutes of time in class to complete an in-depth exam wrapper. This metacognitive tool required students to evaluate and elaborate on their performance with respect to structured reflection including but not limited to, use of available resources, plans for future learning, identification of the highs and lows of student study strategies/efforts. After graded exams were returned, an additional ~5 minutes were provided for students to complete remaining wrapper questions before handing it to the instructor. Exam wrappers were returned to students by the next class period, allowing time for instructors to view the assignment. The data from the exam wrappers were used as a guide for instructors during individual student meetings on growth strategies.

Instructor talk

Course instructors utilized non-biochemistry content language and written feedback. This positively phrased instructor talk included GM messaging, establishing classroom culture, sharing personal experiences, and explaining pedagogical choices [3].

**Supporting Information**

**Figure S1.** Reflection Prompts

**<u>Reflection Prompt #1:</u>** The path to your dream goal is never going to be simple. For this writing reflection, project yourself, 20 or 30 years into the future and write me a letter about your journey (or journeys) to reach that goal. This may include the "highs" and "lows" of that journey and how you overcame obstacles on that path or diverted to create new paths.

**<u>Reflection Prompt #2:</u>** Concept maps are a visual way to draw connections between different concepts/ideas and outline the path or progress towards your understanding of biochemistry. A number of studies from students of various levels have demonstrated the utility in using concept maps as a way to help students efficiently and successfully learn difficult subject material. For this "writing" assignment, I want you to reflect on the concepts you have learned thus far in this course and use that knowledge to construct a concept map containing key points from chapters 2-4. (Think of this as a visual outline).

**<u>Reflection Prompt #3:</u>** Concept maps are a visual way to draw connections between different concepts/ideas and outline the path or progress towards your understanding of biochemistry. A number of studies from students of various levels have demonstrated the utility in using concept maps as a way to help students efficiently and successfully learn difficult subject material. For this "writing" assignment, I want you to reflect on the concepts you have learned thus far in this course and use that knowledge to construct a concept map ***containing in depth details*** from chapter 7 Kinetics and Regulation. (Think of this as a visual outline).

**<u>Reflection Prompt #4:</u>** We have discussed a number of situations whereby an organism can alter or adapt their physiology to suit new environments or purposes. For example, the Sherpa people in the Himalayas can adapt their behavior to function in low oxygen environments, or adjusting our own diet can improve cognitive function through alterations in brain biochemistry etc.

For this reflective assignment, tell me **A)** how have you adapted yourself (whether it be behavior, way of thinking, etc.) to overcome an obstacle to reach a goal in your own life? **B)** How can you use (apply) this experience to improve yourself in other areas? Give a concrete example. **C)** Finally summarize how you overcame that obstacle in <u>Part A</u> in **one BOLDED sentence**. For full credit, please be sure to answer all parts.

**<u>Reflection Prompt #5:</u>** Concept maps are a **visual way** to draw connections between different concepts/ideas and outline the path or progress towards your understanding of biochemistry. A number of studies from students of various levels have demonstrated the utility in using concept maps as a way to help students efficiently and successfully learn difficult subject material. For this "writing" assignment, I want you to reflect on the concepts you have learned thus far in this course and use that knowledge to construct a concept map containing key points from **ONE** of the following chapters 9-12. (Think of this as a visual outline).

**<u>Reflection Prompt #6:</u>** Concept maps are a **visual way** to draw connections between different concepts/ideas and outline the path or progress towards your understanding of biochemistry. A number of studies from students of various levels have demonstrated the utility in using concept maps as a way to help students efficiently and successfully learn difficult subject material. For this FINAL “writing” assignment, I want you to reflect on the concepts you have learned thus far in this course and use that knowledge to construct a concept map containing key points from **ONE** of the following chapters 15-19. (Think of this as a visual outline). You have been doing a fantastic job on these concept maps- end the semester strong!

Figure S2. Example Growth Mindset Messaging.

### Learning is a dynamic process

Biles and Phelps have "grit"

Scientists have demonstrated that *grit is a better predictor of academic success than traditional measures (g.p.a., standardized test scores)* Strayhorn, 2013.

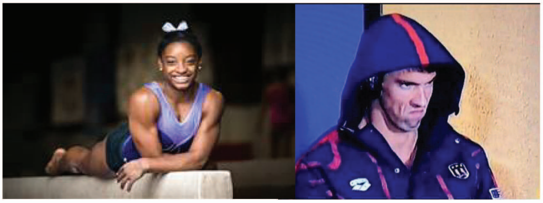

This course is a marathon not a sprint!!

### The Brain Can Change

#### Temporal and Spatial Dynamics of Brain Structure Changes during Extensive Learning

Bogdan Draganski[1], Christian Gaser[2], Gerd Kempermann[3], H. Georg Kuhn[4], Jürgen Winkler[1], Christian Büchel[5], and Arne May[1,5]

The current view regarding human long-term memory as an active process of encoding and retrieval includes a highly specific learning-induced functional plasticity in a network of multiple memory systems. Voxel-based morphometry was used to detect possible structural brain changes associated with learning. Magnetic resonance images were obtained at three different time points while medical students learned for their medical examination. During the learning period, the gray matter increased significantly in the posterior and lateral parietal cortex bilaterally. These structural changes did not change significantly toward the third scan during the semester break 3 months after the exam. The posterior hippocampus showed a different pattern over time: the initial increase in gray matter during the learning period was even more pronounced toward the third time point. These results indicate that the acquisition of a great amount of highly abstract information may be related to a particular pattern of structural gray matter changes in particular brain areas.

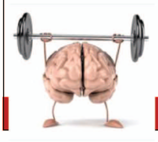

J Neuro 26,(23): 6314-7 (2006)

Figure S3. Example of Student Concept Map

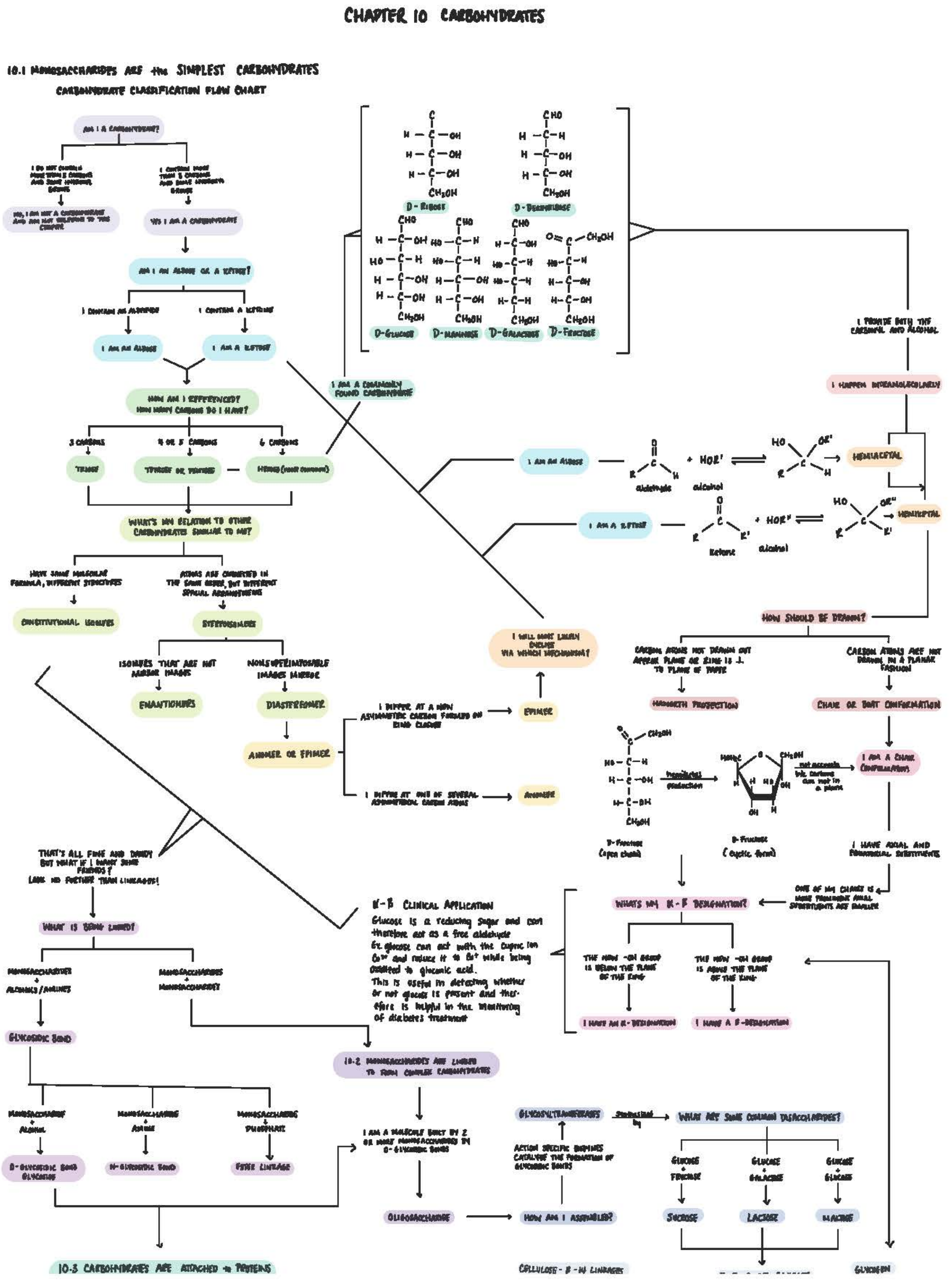

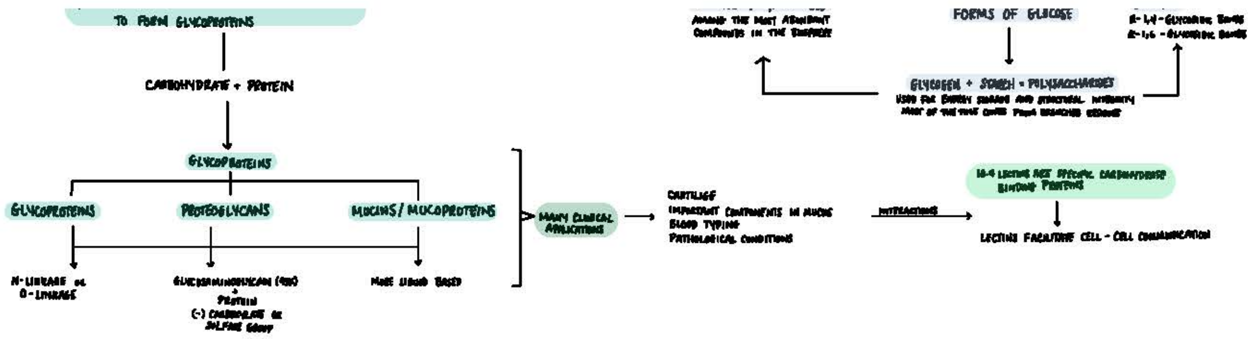

TO FORM GLYCOPROTEINS
CARBOHYDRATE + PROTEIN
GLYCOPROTEINS
GLYCOPROTEINS
PROTEOGLYCANS
MUCINS / MUCOPROTEINS
N-LINKAGE or O-LINKAGE
+
PROTEIN
(-) CARBOXYLATE or SULFATE GROUP
MANY CLINICAL APPLICATIONS
CARTILAGE
BLOOD TYPING
PATHOLOGICAL CONDITIONS
INTERACTIONS
BINDING PROTEINS
FORMS OF GLUCOSE
AMONG THE MOST ABUNDANT COMPOUNDS IN THE BIOSPHERE

**Supporting Information**

**Figure S4.** Growth minded exam wrapper.

<u>Exam Wrapper</u>

Please answer questions **honestly**. You are not assessed on your response, just that you honestly responded! This is an opportunity to identify weaknesses and strengths for your future improvement.

| Question | Yes | Sort of | No | Comments |
|---|---|---|---|---|
| Are there particularly challenging questions that I am proud for how I answered? | | | | |
| What did I do differently in preparing for this exam than the last exam(s)? | | | | |
| Did I set and maintain high standards for myself? | | | | |
| Did I spend enough time to do quality work? | | | | |
| Did I regulate my procrastination, distractions, and temptations in order to complete my work? | | | | |
| Did I make good use of available resources? | | | | |
| Did I ask questions if I needed help? | | | | |
| Did I review and re-review my work for possible errors? | | | | |
| Did I meet with a tutor and/or attend Biochemistry Learning Lab? | | | | |

| |
|---|
| What is one positive thing I did on this exam? (please give a written response in "comments" section. |
